## Supplementary material for "T cell receptor convergence is an indicator of antigen-specific T cell response in cancer immunotherapies"

### Figure 1—figure supplement 1

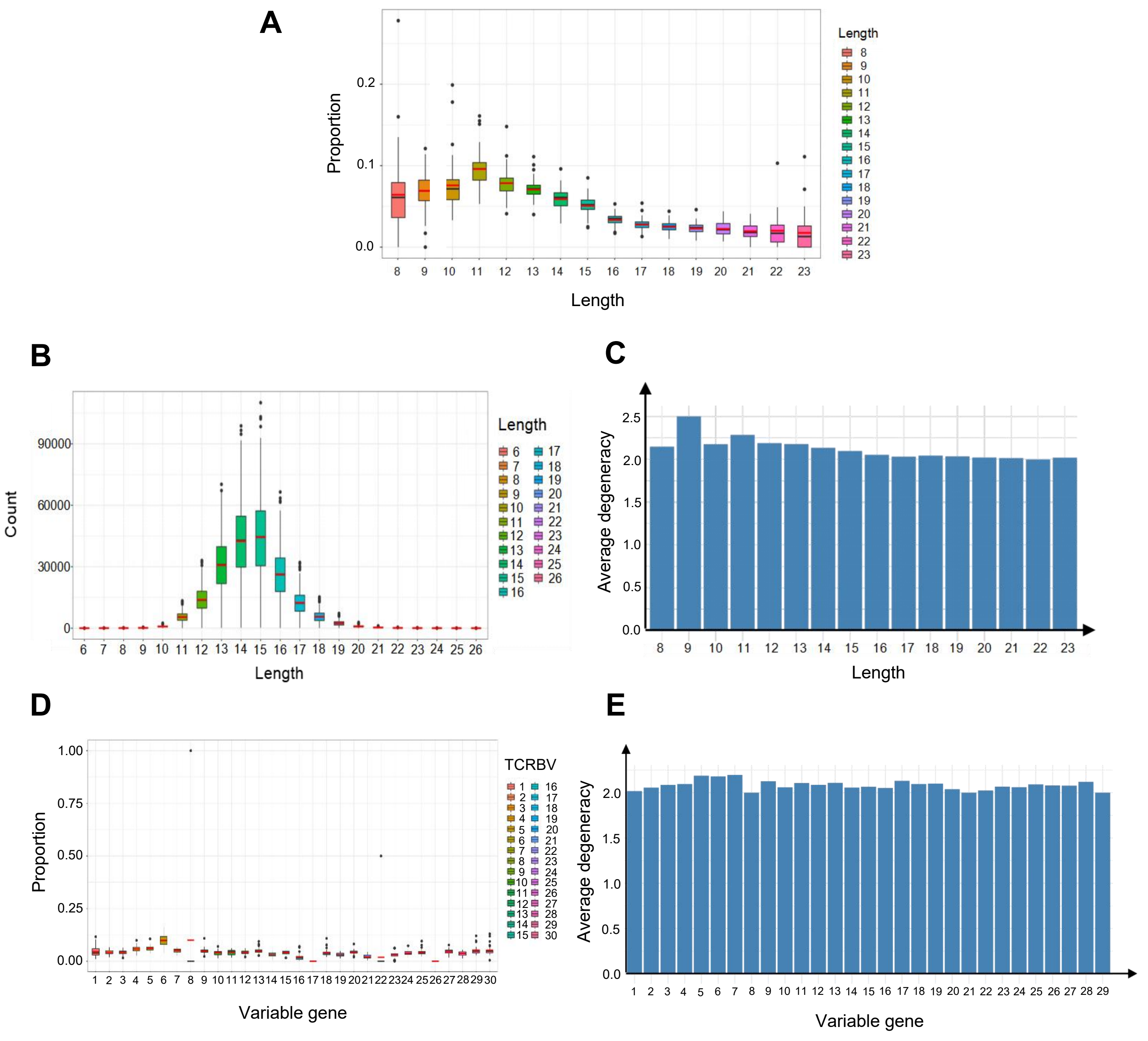

### Figure 3—figure supplement 1

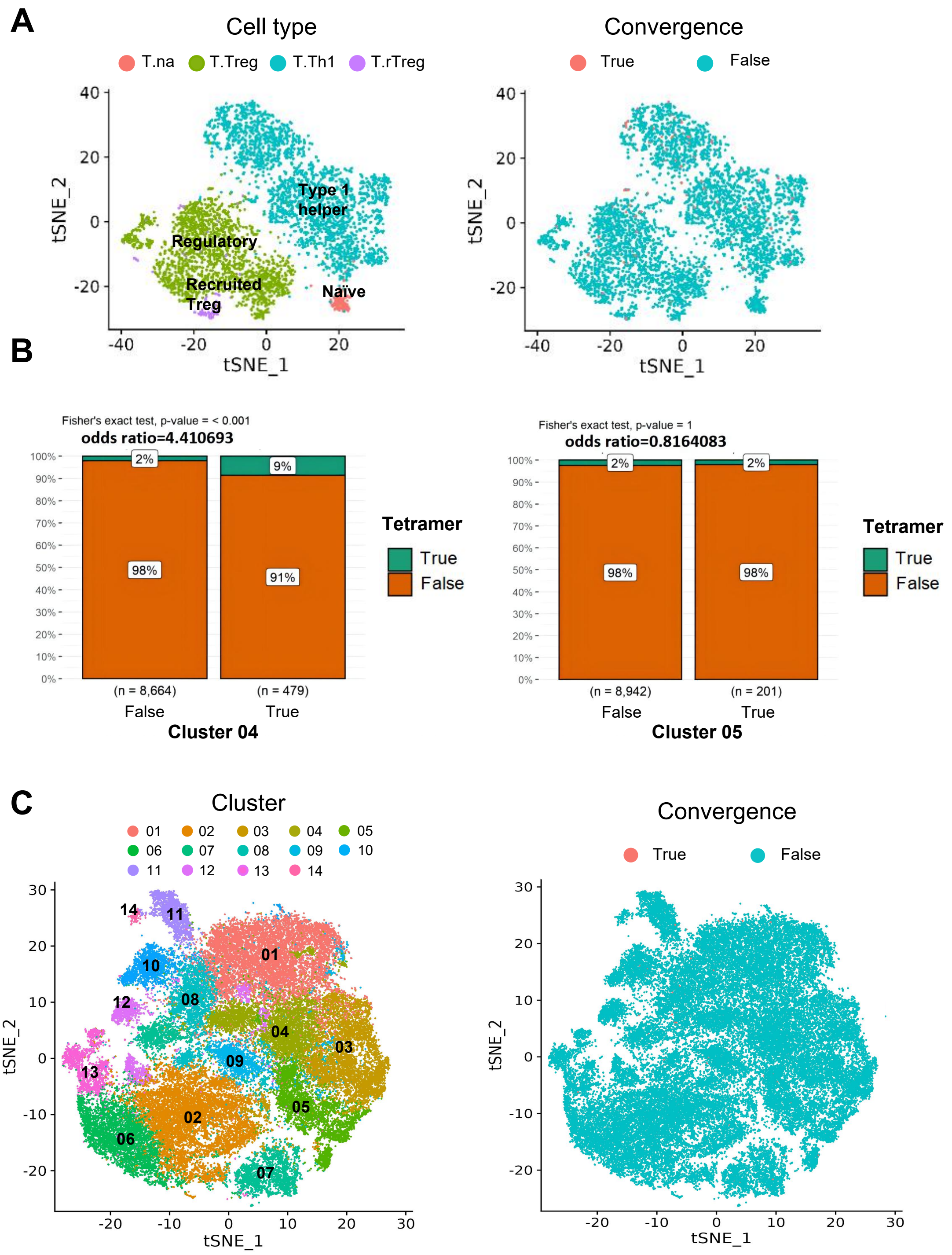

### Figure 3—figure supplement 2

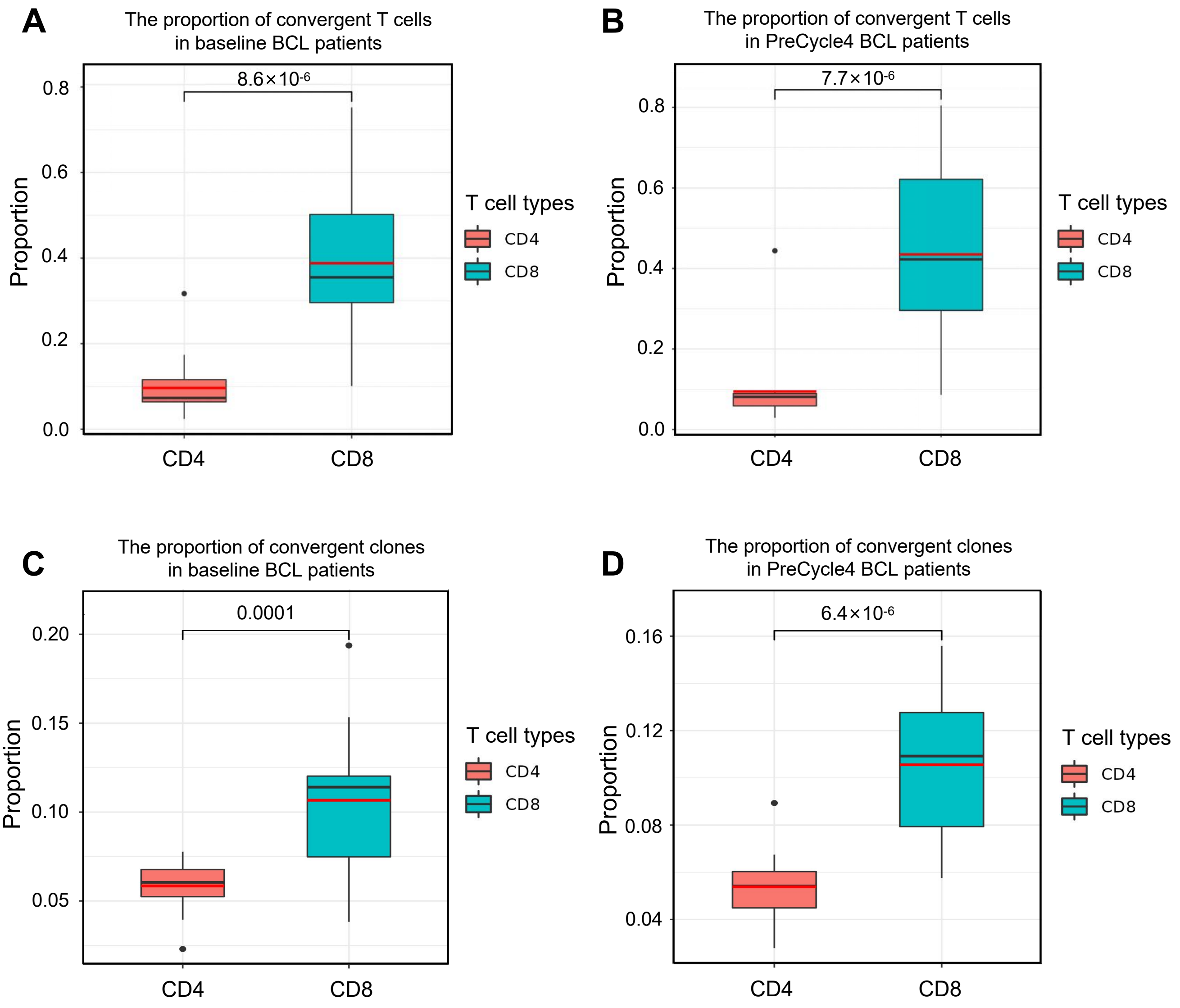

### Figure 4—figure supplement 1

**A**

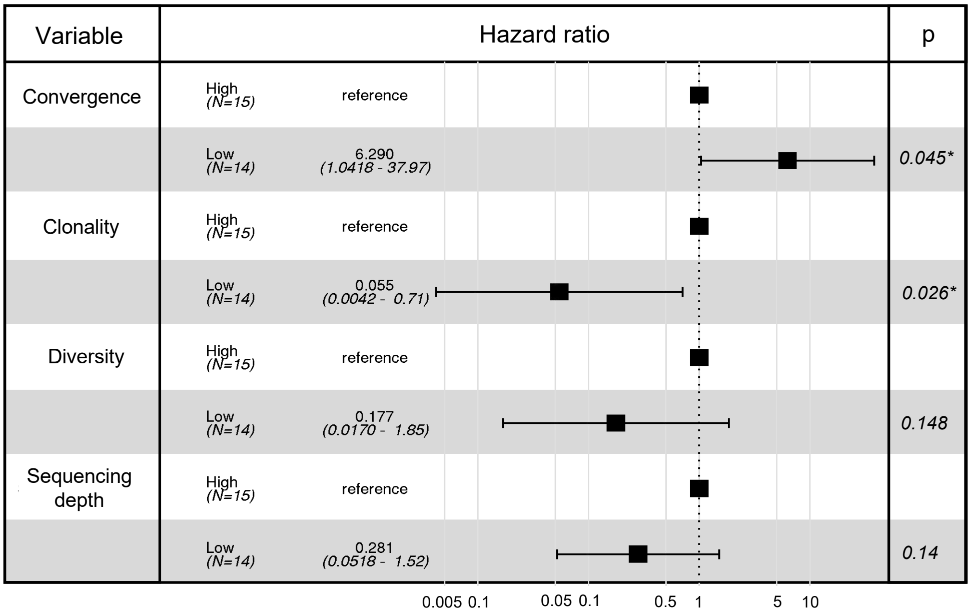


**B**

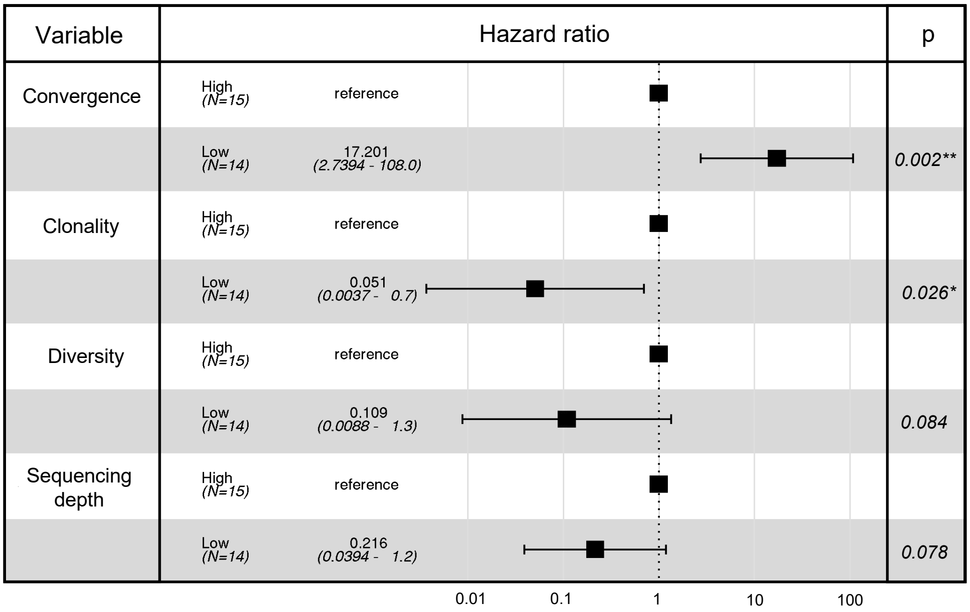

**Supplementary Table 1 Details of datasets used in this study**

| Data Type | Project Name | Researchers | Data Resource |
|---|---|---|---|
| single cell immune profiling | A new way of exploring immunity: linking highly multiplexed antigen recognition to immune repertoire and phenotype | 10x Genomics | 10x Genomicswebsite: https://www.10xgenomics.com/resources/datasets/cd-8-plus-t-cells-of-healthy-donor-1-1-standard-3-0-2 |
| | Concurrent delivery of immune checkpoint blockade modulates T cell dynamics to enhance neoantigen vaccine-generated antitumor immunity | Bo Li; Hongyi Zhang; Longchao | In-house data; GEO accession number: GSE178881 |
| | Pan-cancer single-cell landscape of tumor-infiltrating T cells | S Qin, W Si, A Wang,et al. | GEO accession number: GSE156728 |
| bulk TCRβ-seq | Immunosequencing identifies signatures of cytomegalovirus exposure history and HLA-mediated effects on the T-cell repertoire | Emerson, R., DeWitt, W., Vignali, M. et al. | immuneACCESSDOI：https://doi.org/10.21417/B7001Z |
| | Comprehensive T cell repertoire characterizationof non-small cell lung cancer | Reuben, A., Zhang, J., Chiou, SH. et al. | immuneACCESSDOI：https://doi.org/10.21417/AR2019NC |
| | Tumor and Microenvironment Evolution during Immunotherapy with Nivolumab | N. Riaz, J. J. Havel, V. Makarov, et al. | DOI:https://doi.org/10.1016/j.cell.2017.09.028 |
| | Radiation induces dynamic changes to the T cell repertoire in renal cell carcinoma patients | Chow J , Hoffend NC , Scott AI, et al. | immuneACCESSDOI：https://doi.org/10.21417/JC2020PNAS |
| | Bulk TCRβ-seq of patients with renal cancers | Beshnova et al | https://zenodo.org/record/3894880#.YpwDFy-B2bE |
| | Bulk TCRβ-seq of patients with high-Grade ovarian cancer | Beshnova et al | https://zenodo.org/record/3894880#.YpwDFy-B2bE |
| | A large-scale database of T-cell receptor beta (TCRb) sequences and binding associations from natural and synthetic exposure to SARS-CoV-2 | Nolan S, Vignali M, Klinger M. et al. | immuneACCESS DOIhttps://doi.org/10.21417/ADPT2020COVID |
| | Contribution of systemic and somatic factors to clinical response and resistance to PD-L1 blockade in urothelial cancer: An exploratory multi-omic analysis | Snyder A, Nathanson T, Funt SA. et al. | immuneACCESS DOIhttps://doi.org/10.21417/B7MG68 |
| | Association of Tumor Microenvironment T-Cell Repertoire and Mutational Load With Clinical Outcome After Sequential Checkpoint Blockade in Melanoma | Erik Y, Marissa V, Richard KW. et al. | immuneACCESS DOIhttps://doi.org/10.21417/EY2019CIR |
| | A peripheral immune signature of responsiveness to PD-1 blockade in patients with classical Hodgkin lymphoma | Cader, F.Z., Hu, X., Goh, W.L. et al. | immuneACCESSDOI：https://doi.org/10.21417/FZC2020NM |

**Supplementary Table 2 The clusters annotation in human pan-cancer data**

| Clusters | Defined Cell Types | Signature Genes |
|---|---|---|
| CD4-01-FOS | $CD4^+$ Effector memory T cells | FOS, CCL4, CD69, IL7R, KLRB1 |
| CD4-02-FOXP3 | $CD4^+$ Tregs | FOXP3, CTLA4, IKZF2, IL2RA |
| CD4-03-CCR7 | $CD4^+$ Naive T cells | CCR7, SELL, LEF1, S1PR1 |
| CD4-04-CCR6 | $CD4^+$ effector T cells | CCR6, KLRB1, IL7R, S100A11 |
| CD4-05-GZMA | $CD4^+$ activating T cells | GIMAP7, RPS29, RPS21 |
| CD4-06-IL2RA | $CD4^+$ Tregs | FOXP3, IL2RA, CTLA4, LAG3, IKZF2 |
| CD4-07-CD200 | $CD4^+$ Follicular helper T cells | CD200, TOX, CXCR5, TNFSF8 |
| CD4-08-S100A11 | $CD4^+$ effector T cells | GZMA, S100A11, KLRB1, ITGB1, CXCR6 |
| CD4-09-CD69 | $CD4^+$ effector T cells | CD69, IFNG, TNF |
| CD4-10-IFNG | $CD4^+$ effector T cells | IFNG, NKG7, CCL4, CST7, PRF5, GZMA, GZMB, CCL3 |
| CD4-11-NKG7 | $CD4^+$ effector T cells | NKG7, CCL4, CST7, PRF5, GZMA, GZMB, TBX21 |
| CD4-12-ISG15 | $CD4^+$ IFNs responsive T cells | ISG15, IFI6, MX1, IFIT1, LY6E |
| CD4-13-STMN1 | $CD4^+$ proliferative T cells | STMN1, TUBB, TUBA1B, MKI67 |
| CD4-14-GNLY | $CD4^+$ effector T cells | GNLY, NKG7, PRF1, CCL4, KLRB1, GZMB, GZMA, ZNF683 |
| CD8-01-XCL1 | $CD8^+$ resident memory T cells | XCL1, CXCR6, GZMB, ITGAE, ITGA1 |
| CD8-02-GZMK | $CD8^+$ effector memory T cells | GZMK, CD28, EOMES, CCR7, CXCR3, CXCR5 |
| CD8-03-TIGIT | $CD8^+$ exhausted T cells | TIGIT, CTLA4, LAYN, ENTPD1, LAG3, HAVCR2, TOX, PDCD1 |
| CD8-04-ZNF683 | $CD8^+$ effector memory T cells | ZNF683, CD27, GZMA, EOMES, CXCR3 |
| CD8-05-FGFBP2 | $CD8^+$ cytotoxic T cells | FGFBP2, GNLY, NKG7, PRF1, GZMB |
| CD8-06-IFNG | $CD8^+$ cytotoxic T cells | CCL4, IFNG, TNF, CD69, CCL3 |
| CD8-07-IHSPA6 | $CD8^+$ dysfunctional T cells | HSPA6, HSPA1A, HSPA1B, HSPH1 |
| CD8-08-PRF1 | $CD8^+$ cytotoxic T cells | NKG7, PRF1, GZMB, CCL3 |
| CD8-09-IL7R | $CD8^+$ naive T cells | CCR7, IL7R, SELL, S1PR1, LEF1, CD28 |
| CD8-10-IFI6 | $CD8^+$ IFNs responsive T cells | IFI6, MX1, IFI27, IFIT1, STAT1 |
| CD8-11-STMN1 | $CD8^+$ proliferative T cells | STMN1, TUBB, TUBA1B, MKI67 |
| CD8-12-KLRB1 | $CD8^+$ cytotoxic T cells | KLRB1, CCR6, TNF, RORC, FOS |
| CD8-13-CCR7 | $CD8^+$ naive T cells | CCR7, SELL, LEF1, IL7R, S1PR1. SELL |
| CD8-14-CXCR6 | $CD8^+$ resident memory T cells | CXCR6, PDCD1, ITGAE, LAYN |
| CD8-15-CTLA4 | $CD8^+$ exhausted T cells | CTLA4, LAYN, ENTPD1, TIGIT, LAG3 |
